# Ventricle stimulation as a potential gold-standard control stimulation site for transcranial focused ultrasound stimulation

**DOI:** 10.1101/2024.05.29.596387

**Authors:** Cyril Atkinson-Clement, Marcus Kaiser, Matthew A. Lambon Ralph, JeYoung Jung

## Abstract

This research investigates whether ventricular-focused ultrasound stimulation (ventricle-FUS) can serve as an effective control in studies using transcranial FUS, a non-invasive technology for brain modulation. FUS has notable potential for therapeutic applications but requires a robust control to accurately assess its effects. We evaluated the effectiveness of ventricle-FUS, as an active, non-cerebrum control for FUS research, comparing it to sham stimulation. We conducted a comprehensive assessment of ventricle-FUS, employing both questionnaires and multiple neuroimaging metrics such as grey matter and white matter volumes, and functional connectivity. Ventricle-FUS did not alter any of these metrics, thereby successfully retaining the auditory, somatosensory, and experiential elements of FUS without affecting brain structure or function. Importantly, participants were unable to distinguish whether they received ventricle-FUS or sham FUS. Our findings confirm that ventricle-FUS establishes it as a reliable control approach in FUS research, crucial for advancing its therapeutic applications.

## Introduction

Low-intensity transcranial focused ultrasound stimulation (FUS) is a very promising methods for treating brain disorders (*1*). It consists of applying acoustic energy to a specific part of the brain to transiently change its functioning. Unlike others non-invasive brain stimulation (NIBS) techniques, FUS has many concurrent advantages: it is safe (*2*), painless, spatially accurate (*3*), can reach both cortical and deep territories, and can increase or decrease brain activity depending on the used parameters (*4*).

However, FUS is still under development and several methodological considerations must be addressed before going further. Among them is the need for a true, validated sham condition, especially because FUS generates acoustic features (a sound is produced during the stimulation), tactile feelings (contact between the transducer with coupled gel and the head, and vibrations) and experiential elements (the procedures and experiences of being stimulated). The ITRUSST Consortium (International Transcranial Ultrasonic Stimulation Safety and Standards; https://itrusst.com/index.htm) already referenced six different possible sham conditions ranked as follows: 1) no stimulation (but with the transducer over the participant’s head); 2) flip-over the transducer (could damage the transducer); 3) blocking ultrasound waves with a cap or using a high impedance between the transducer and the head (could damage the transducer); 4) using an ineffective protocol; 5) defocusing the acoustic waves; 6) using an inactive control. However, most of the published study only used a sham condition of level 1 or 2 (see Table.1).

**Table 1.** List of the control condition used in published human’s studies (only those with a sham condition)

| Control type (levels 1-6) | References |
| --- | --- |
| (1) No stimulation – transducer over the head | Hameroff et al., 2013, Brain Stim (5)<br>Lee et al., 2015, Sci rep (6)<br>Lee et al., 2016, BMC (7)<br>Lee et al., 2016, Sci rep (8)<br>Ai et al., 2018, BMC Neurosci (9)<br>Gibson et al., 2018, Front Neurol (10)<br>Badran et al., 2020, Brain Stim (11)<br>Braun et al., 2020, Brain Stim (12)<br>Fomenko et al., 2020, eLife (13)<br>Reznik et al., 2020, Neurol Psy Brain Res (14)<br>Sanguinetti et al., 2020, Front Hum Neurosci (15)<br>Schimek et al., 2020, Front Human Neurosci (16)<br>Guerra et al., 2021, Brain Sci (17)<br>Johnstone et al., 2021, Brain Stim (18)<br>Xia et al., 2021, NeuroImage (19)<br>Heimbuch et al., 2022, Plos One (20)<br>Park et al., 2022, Comp Meth Prog Bio (21)<br>Zhang et al., 2021, iScience (22)<br>Kim et al., 2023, Plos one (23)<br>Yaakub et al., 2023, Nat com (24)<br>Chou et al., 2024, Brain Stim (25) |
|  | Legon et al., 2024, Pain (26)<br>Peng et al., 2024, Front Hum Neurosci (27)<br>Riis et al., 2024, J Neural Eng (28)<br>Strohman et al., 2024, J Neurosci (29) |
| (2) Flip-over | Legon et al., 2014, Nat Neurosci (30)<br>Mueller et al., 2014, Brain Stim (31)<br>Legon et al., 2018, Sci rep (32)<br>Fomenko et al., 2020, eLife (13)<br>Xia et al., 2021, NeuroImage (19)<br>Samuel et al., 2022, Brain Stim (33)<br>Zeng et al., 2022, Annals of Neurol (34)<br>Fine et al., 2023, eLife (35)<br>Ziebell et al., 2023, Brain Stim (36) |
| (3) Blocking cap<br>(3) High impedance | Beisteiner et al., 2020, Adv sci (37)<br>Legon et al., 2018, Hum Brain Mapp (38)<br>Liu et al., 2021, Ultra Med Biol (39)<br>Yu et al., 2021, IEEE Trans BioMed Eng (40) |
| (4) Ineffective protocol | - |
| (5) Defocusing | - |
| (6) Inactive control | - |

In contrast, transcranial magnetic stimulation (TMS), a well-established non-invasive brain stimulation (NIBS) technique, typically employs a control stimulation technique where active stimulation is delivered to a task-irrelevant brain region, which acts as a control site (commonly the vertex). This approach retains and thus controls for the sensory effects, such as the ‘click’ sound and the tactile sensations caused by the magnetic field of the TMS coil. Traditionally, the vertex located at the top of the head, corresponding to Cz in the international 10-20 system (*41*), has been frequently used as the control site in TMS research, often referred to as the ‘Empty Quarter’ (*42*). Many studies have shown that stimulating the vertex does not result in behavioural changes (*43-47*). Unlike other forms of NIBS, FUS has the ability to determine not only the focus but also the depth of stimulation. This opens up the novel opportunity for a novel, effective and active FUS control: stimulation of the cerebrospinal fluid (CSF) in the posterior lateral ventricle as a control site.

The aim of this study was to develop and validate a gold-standard control stimulation for FUS studies. We investigated how ventricle stimulation affects participants’ reported experiences (see Table S1) as well as brain structure and function using structural imaging and resting-state fMRI (rs-fMRI). Participants received theta burst FUS (tbFUS) protocol(*34*) at the left lateral ventricle for the ventricle stimulation while we played simulated tbFUS sound via a bond-conducted headset during sham stimulation (Fig. 1). Then, we acquired the structural and resting-state functional magnetic resonance (rs-fMRI) imaging (Fig. 1). We compared two control conditions: (1) a typical sham (no stimulation); and (2) active stimulation of the left posterior lateral ventricle. Specifically, we examined changes in grey matter (GM), white matter (WM) and CSF around the ventricle, as well as rs-fMRI measures including fractional amplitude of low-frequency fluctuations (fALFF), regional homogeneity (ReHo) and eigenvector centrality. Previous examinations of ‘control’ TMS to the vertex have observed decreased blood-oxygen-level-dependent (BOLD) signal changes in the default mode network (DMN) even though there were no changes in overall brain activity (*48*). Accordingly, we also investigated functional connectivity changes within seven well-established functional brain networks: DMN, frontoparietal network (FPN), salience network (SN), dorsal attention network (DAN), sensorimotor network (SMN), visual network (VN), and auditory network (AN). We expected that targeting the lateral ventricle would be an effective active control condition since it should induce no brain changes (no difference with the no stimulation condition) yet retains the sensory and experiential phenomena related to brain-targeted FUS.

**Figure 1.**
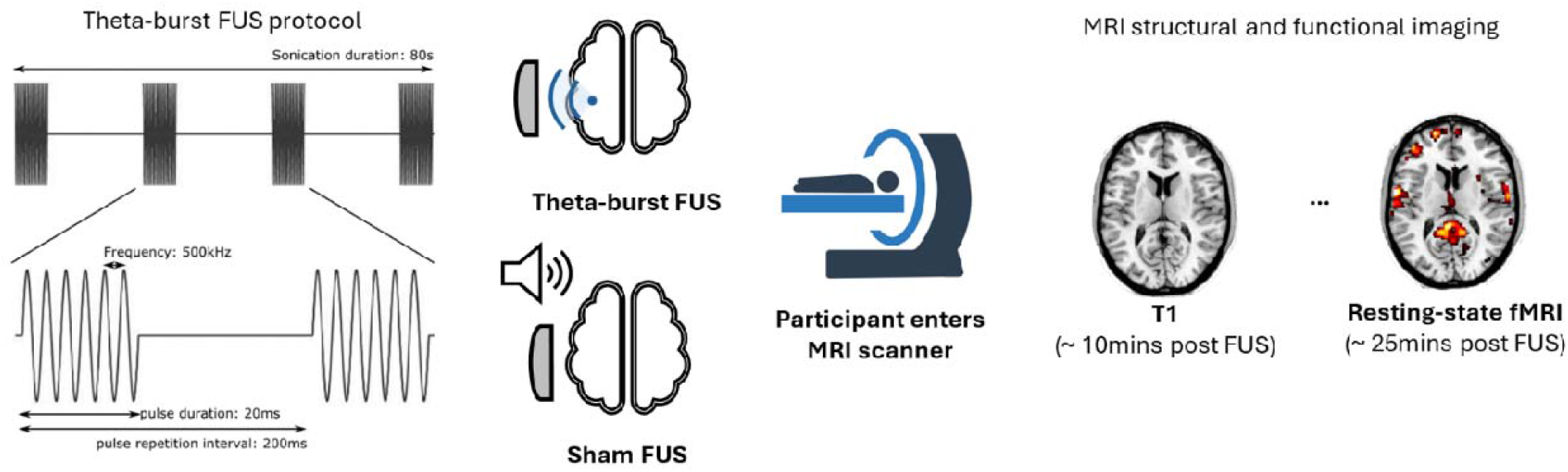
theta-burst FUS protocol (left) and experimental procedure applying theta-burst FUS to the lateral ventricle or applying no FUS (sham), followed by an imaging session (right).

## Results

We successfully stimulated the left lateral ventricle using FUS. Fig. 2 shows the activated volumes for each participant during the ventricle stimulation.

**Figure 2.**
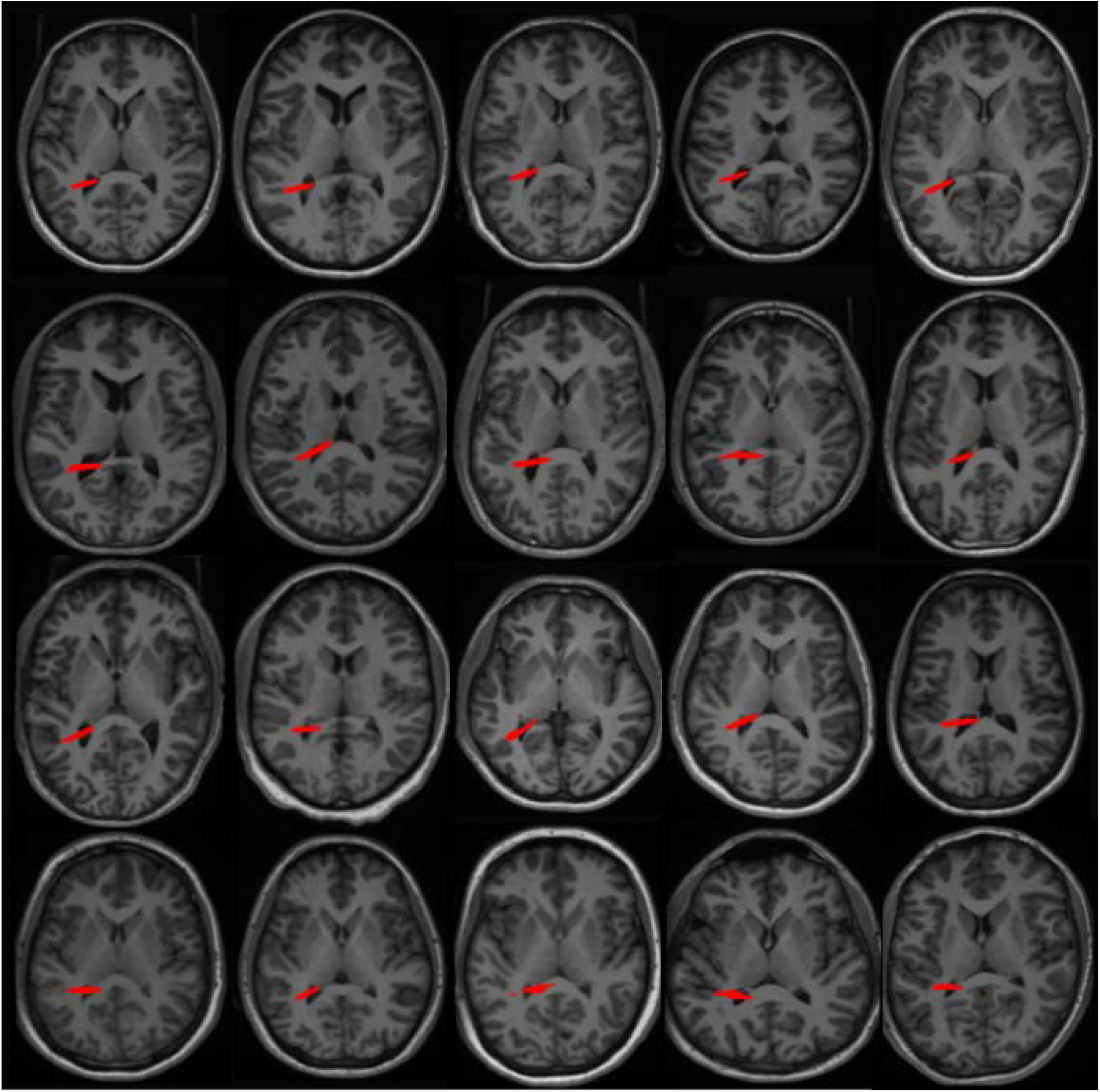
Activated volumes for the 20 participants during the lateral ventricle stimulation, according to k-plan acoustic simulations.

### Questionnaires results

No participants reported experiencing any discomfort during the experiment. Participants provided feedback on their experiences during both the lateral ventricle FUS and sham sessions. All participants reported no discomfort linked to FUS. Analysis of the questionnaire responses revealed no significant differences as shown in Table 2. Participants who experienced ‘unusual sensations on the skin’ or ‘tingling’ attributed these feelings to the ultrasound gel. Those who reported ‘neck pain’ attributed it to the chair used during the session. Reports of ‘difficulty paying attention’ and ‘sleepiness’ were associated with the lengthy duration of the session. Lastly, any incidents of ‘dizziness’ were linked to MRI scanning, not the FUS. Approximately 70% of participants reported additional symptoms, yet all noted ‘hearing a faint clicking’ or ‘tapping sound’ during both the sham and ventricle stimulation sessions. In addition, there were no significant differences between the ventricle and ATL sessions (see Table S2). None of the participants were able to distinguish between the sham, active-ventricle, and active ATL sessions.

**Table 2.** The results of questionnaires. The number was the number of participants.

| | Sham FUS | Ventricle FUS | $\chi^2$ | p |
| --- | --- | --- | --- | --- |
| Headache | 1 | 0 | 1.723 | 0.422 |
| Unusual feeling on the skin | 0 | 2 | 3.648 | 0.161 |
| Neck pain | 1 | 2 | 2.135 | 0.344 |
| Tingling | 3 | 3 | 0.069 | 0.966 |
| Itchiness | 0 | 0 | – | – |
| Difficulty paying attention | 4 | 3 | 0.401 | 0.818 |
| Unusual feelings, attitude, emotions | 1 | 0 | 1.723 | 0.422 |
| Sleepiness | 5 | 5 | 3.425 | 0.489 |
| Change in hearing | 1 | 1 | 0.928 | 0.629 |
| Nausea/stick to stomach | 0 | 0 | – | – |
| Dizziness | 2 | 2 | 1.919 | 0.383 |
| Anxious/worried/nervous | 3 | 0 | 5.346 | 0.069 |
| Forgetful | 0 | 0 | – | – |
| Difficulty with your balance | 0 | 1 | 1.221 | 0.543 |
| Other |  |  |  |  |
| (heard faint clicking/tapping sound during the stimulation) | 13 | 14 | 3.709 | 0.157 |

### Structural results

To examine the effects of ventricle FUS on the brain structure, we performed voxel-based morphometry (VBM) analysis. The whole brain analysis revealed no significant differences between ventricle stimulation and sham stimulation. The lateral ventricle ROI analysis also demonstrated no significant changes on the GM volume (t = 1.02, p = 0.322; BF10 = 0.366), WM volume (t = -0.487, p = 0.632; BF10 = 0.258), and CSF (t = 0.591, p = 0.562; BF10 = 0.272) between the ventricle and sham stimulations. The results are illustrated in Fig. 3.

**Figure 3.**
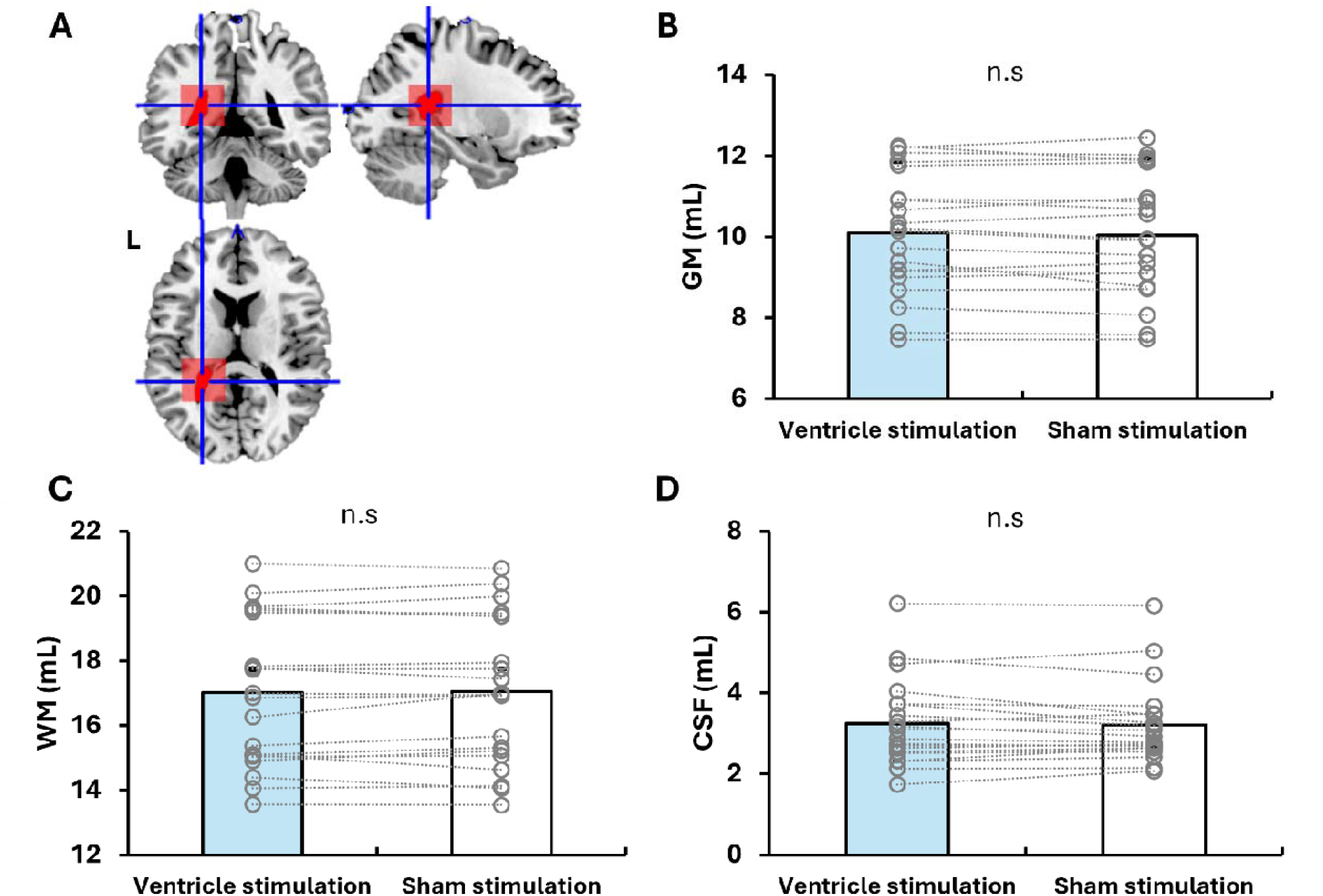
Structural differences between sham and ventricle stimulations. **A**. Ventricle region of interest (ROI) in the left hemisphere. **B**. Gray matter (GM) volume results. **C**. White matter (WM) volume results. **D**. Cerebrospinal fluid (CSF) volume results. Each circle denotes an individual participant. Light blue bars represent ventricle stimulation, while white bars depict the sham stimulation.

### Functional results

At the whole-brain level, no significant difference was found between the sham and ventricle conditions on any of the three rs-fMRI metrics (fALFF, ReHo, eigenvector centrality; see Fig. 4).

**Figure 4.**
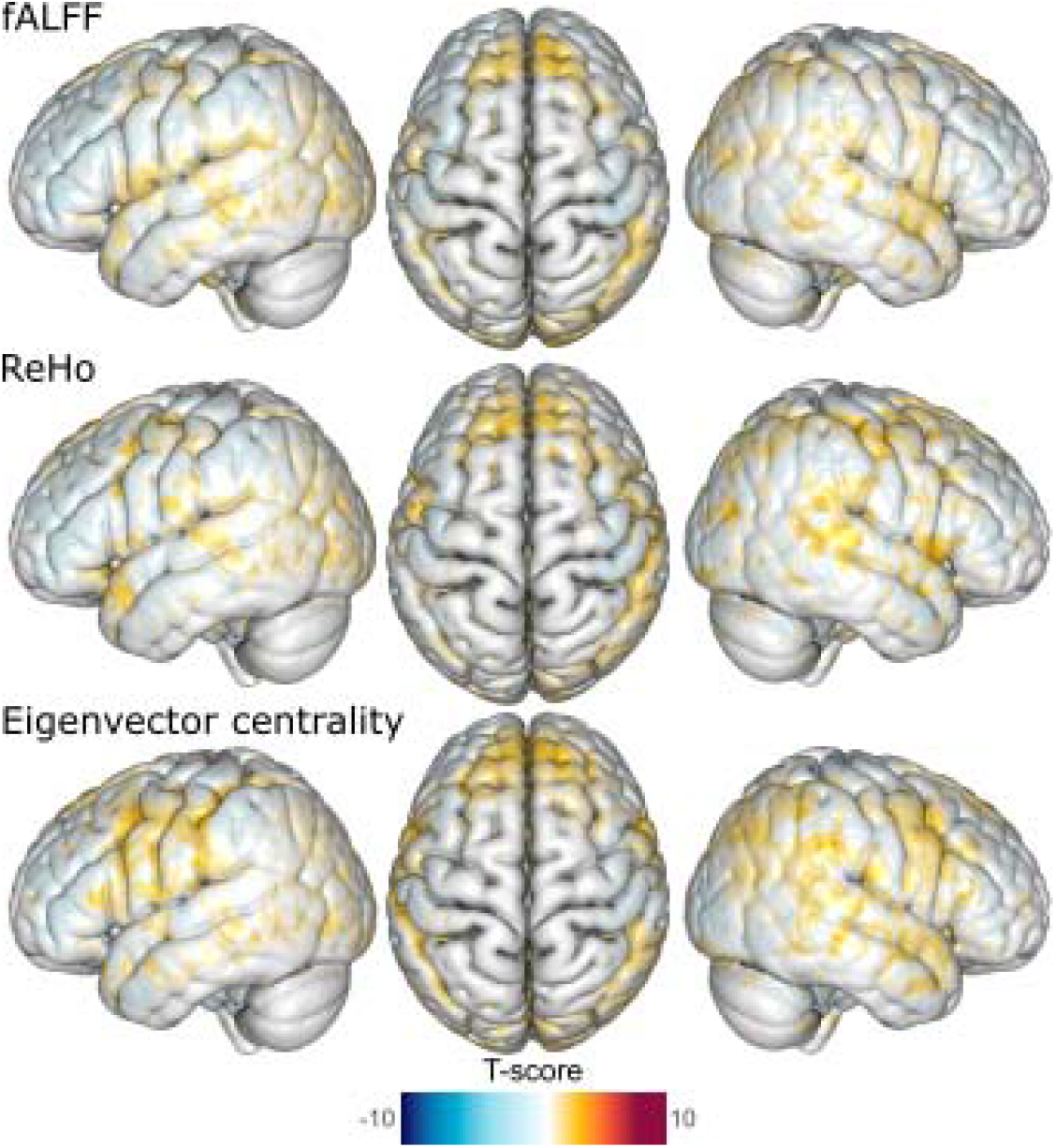
Whole-brain functional differences between sham and ventricle stimulations. No difference reached the threshold for significance.

To investigate the ventricle FUS effects in functional brain networks, we analyzed the FC across seven functional brain networks (Fig. 5). We found no significant FC differences in the DMN (t = 0.119, p = 0.907; BF10 = 0.234), FPN (t = 0.557, p = 0.584; BF10 = 0.267), SN (t = -0.080, p = 0.933; BF10 = 0.233), DAN (t = 0.902, p = 0.378; BF10 = 0.333), SMN (t = 0.343, p = 0.735; BF10 = 0.245), VN (t = -0.898, p = 0.381; BF10 = 0.332), and AN (t = -1.154, p = 0.263; BF10 = 0.416).

**Figure 5.**
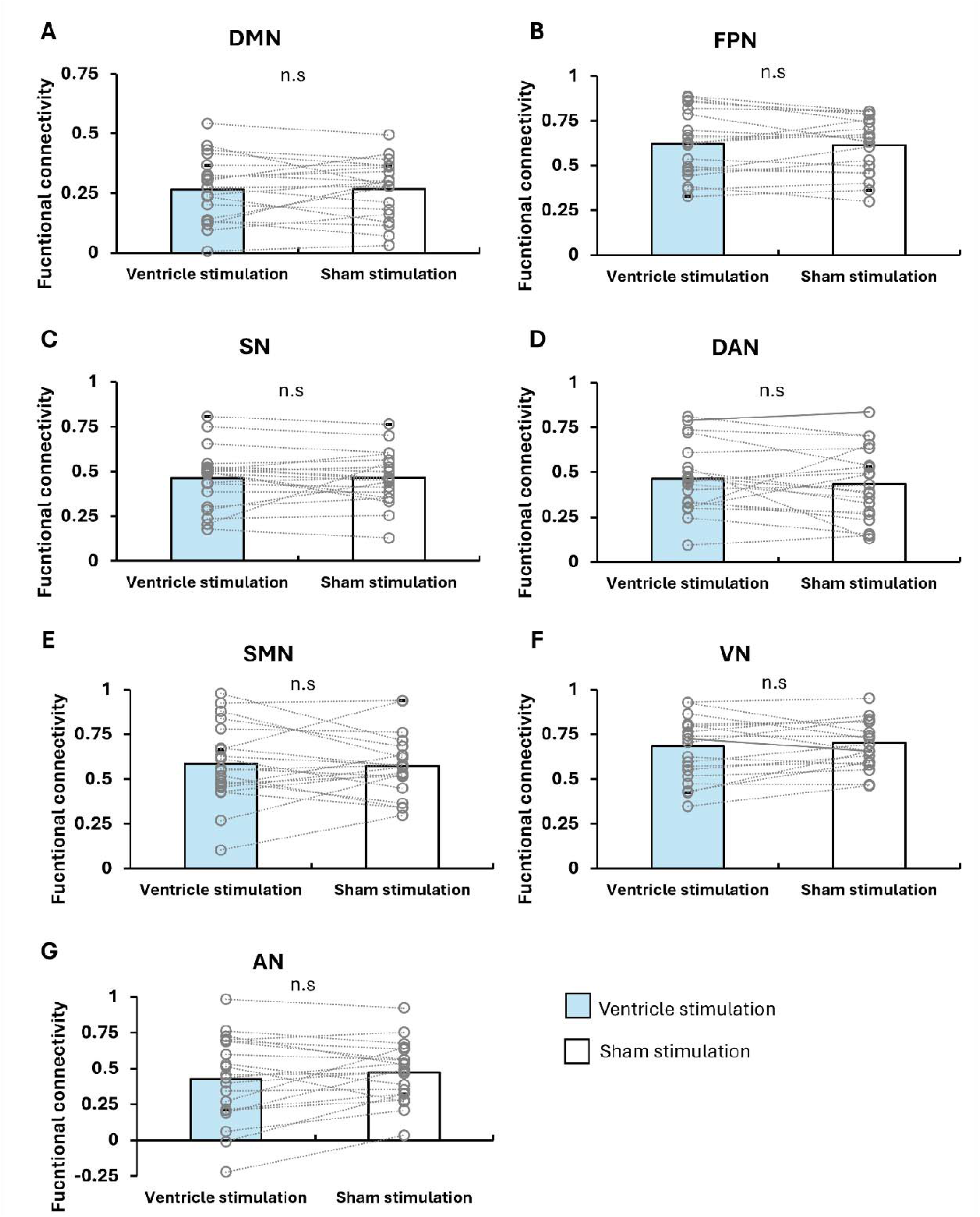
Functional connectivity analysis results across seven brain networks. **A**. Functional Connectivity in the Default Mode Network (DMN), **B**. Functional Connectivity in the Frontoparietal Network (FPN), **C**. Functional Connectivity in the Salience Network (SN), **D**. Functional Connectivity in the Dorsal Attention Network (DAN), **E**. Functional Connectivity in the Sensorimotor Network (SMN), **F**. Functional Connectivity in the Visual Network (VN), **G**. Functional Connectivity in the Auditory Network (AN). Each circle represents an individual participant. Light blue bars indicate ventricle stimulation, while white bars represent sham stimulation.

## Discussion

In this study, we assessed whether ventricular-focused ultrasound (ventricle-FUS) serves as an optimal active control for focused ultrasound stimulation (FUS) studies. Our results strongly support this aim: ventricle-FUS retains the auditory and somatosensory sensory and procedural experiential features associated with actual FUS, without producing measurable changes in brain structure or function. This is particularly significant given that our evaluation spanned multiple metrics, including GM, WM, and CSF densities, as well as functional connectivity, fALFF, ReHo, and eigenvector centrality. We observed no significant alterations in any of these areas, whether analysed at the whole-brain level, using network atlases, or within the lateral ventricle ROI.

When this theta-burst FUS protocol has been applied to the primary motor cortex, it enhances corticospinal excitability, reduces short-interval intracortical inhibition, and increases intracortical facilitation, thereby confirming its facilitatory effects on the motor system (*34*). Importantly, it also improved performance on a visuomotor task post-stimulation. Yaak and colleagues (*24*) employed this protocol to stimulate the anterior cingulate cortex (ACC) and posterior cingulate cortex (PCC), showing increased functional connectivity within the salience network after ACC stimulation and the default mode network after PCC stimulation, with effects persisting for at least 50 minutes post-stimulation. In contrast, our findings revealed no behavioural and functional changes following lateral ventricle FUS. The absence of brain activity changes with ventricle-FUS is notable, especially considering the conservative significance threshold we applied (using uncorrected p-values). This highlights the utility of ventricle-FUS as a valid control that delivers the physical experiences of FUS without the neuromodulatory effects, thus providing a robust method for isolating the neural impacts of FUS.

Importantly, this ventricle-FUS control goes beyond the limitations of previous attempts to develop an effective control condition for FUS studies. These include the primary issue of the audible noise produced during stimulation, which could potentially bias the participants’ perceptions of the experiment. While one study suggested that using headphones to mask this sound might be effective (*12*), another indicated that neither auditory masking with headphones nor the use of ramping parameters adequately obscured the noise generated during FUS (*18*). The ventricle-FUS simply and effectively side-steps this issue given that it is an active FUS protocol.

In conclusion, we demonstrated that the ventricle FUS did not induce any structural and functional changes at any voxels throughout the whole brain and that, experientially, participants were unable to distinguish ventricle FUS from active FUS. Our findings affirm the potential of using the ventricle as a control site in FUS research. This strategy enhances the methodological rigor by ensuring that the observed effects are truly attributable to the brain-targeted stimulation.

## Materials and Methods

### Participants

Twenty healthy adult participants (6 males, mean age = 21.95 ± 3.24 years old ranging from 19 to 33 years old) participated in the study. All participants were right-handed (*49*). They provided informed written consent prior to the study. The study was approved by the ethics committee of School of Psychology, University of Nottingham (F1417).

### Experimental design and procedure

Participants took part in three sessions for the study. In each visit, participants received FUS prior to scanning. Immediately following the FUS, they were transported to the MRI centre for scanning. The first session involved sham FUS, while the subsequent sessions included active FUS targeting either the ventricle or the ventral anterior temporal lobe (ATL). Sessions were a minimum of five days apart, with an average of 16.20 ± 7.96 days between the sham and ventricle sessions. At the end of each session, participants evaluated their experience of FUS using a questionnaire (Supplementary Table 1). The study design and procedure are displayed in Fig. 1. We focus only on the sham and ventricle stimulation sessions in this study.

### Transcranial focused ultrasound stimulation and acoustic simulations

During the ventricle stimulation, FUS was applied with the following theta-burst parameters: central frequency = 500 kHz, pulse duration = 20 ms, pulse repetition interval = 200 ms, duty cycle = 10%, ISPPA = 54.51 W/cm2, total duration = 80sec (Fig. 1). This protocol, developed by Zeng et al (*34*) as a facilitatory protocol, has been shown to induce changes in resting-state functional connectivity in healthy individuals (*24*), and is in line with the safety guidelines (*50*) (U.S. Food and Drug Administration. Marketing Clearance of Diagnostic Ultrasound Systems and Transducers). During the sham stimulation, participants wore a bone-conducting headset that played a simulated theta-burst FUS sound for 80 seconds (Fig. 1).

To ensure that our protocol accurately reached the brain target, we ran acoustic simulations for all participants with k-plan V1.0 (Brainbox Ltd., Cardiff, United Kingdom). The stimulations used both the acquired T1-weighted and a pseudo computed tomography (PCT) scan that was estimated for each participant from the T1-weighted scan (*51*). The activated volume estimation was based on the full width at half maximum (FWHM) of the ultrasound beam (i.e., every voxel with an applied pressure higher than 50% of the maximum was considered as part of the activated volume). Acoustic simulations confirmed that the ventricle was correctly reached for all participants (Figure 2). For the whole group, the simulations reported a peak pressure of 191.9 ± 28.06 kPa, an ISPPA of 1.25 ± 0.4 W/cm2, a mechanical index of 0.27 ± 0.04, a maximal temperature increase of 0.09 ± 0.02 C° and an activated volume of size 0.33 ± 0.13 ml. All these outcomes are far below the safety limits recommended (*52*).

### Magnetic resonance imaging acquisition and preprocessing

Images were acquired using a General Electric (GE) SIGNA Premier 3T MR scanner with a 48-channel head coil (GE Healthcare, USA), with an established procedure (*53*).

T1-weighted images were acquired using 3D MPRAGE sequence (voxel size = 1mm isotropic, field of view [FOV] = 256, matrix = 256, 256 sagittal slices, inverse time [TI] = 800ms, flip angle [FA] = 8°). These images were pre-processed using CAT12 (https://neuro-jena.github.io/cat12-help/#intro) (*54*) implemented in SPM12 (Wellcome Department of Imaging Neuroscience, www.fil.ion.ucl.ac.uk/spm) for voxel-based morphometry (VBM) analysis (*55*).

They were segmented into 6 categories: GM, WM, CSF, and three background types. Then, they were spatially normalised to the SPM’s template using Geodesic Shooting (*56*). GM and WM images were modulated and smoothed with an 6mm full width at half maximum (FWHM) Gaussian kernel.

Rs-fMRI data were collected using whole-brain 2D GE-EPI sequence (TR = 1400ms, TE = 35ms, flip angle = 68°, in-plane FOV = 212 × 212mm, 57 slices, slice thickness = 2 mm, voxel size = 2mm isotropic, hyperband factor = 3, ARC factor =2, 220 volumes, 5.13 min total scan time). Additionally, to correct for echo-planar imaging (EPI) distortions, two SE-EPI images with opposite phase encoding directions will be acquired, sharing the same geometry, echo spacing, and phase encoding direction parameters as the GE fMRI scans.

Functional images were pre-processed using the latest version of the NIHR Nottingham BRC imaging pipeline, as detailed on their GitHub page (https://github.com/SPMIC-UoN) (*57*). This pipeline incorporates and utilizes tools from leading advanced image analysis software such as FSL, ANTs, SPM, and FreeSurfer. Resting-state fMRI data were corrected for EPI distortion, motion, and slice timing, and then underwent normalization and smoothing with a 5mm FWHM. To examine the effects of the lateral ventricle FUS, we extracted metrics of functional connectivity (FC), fractional Amplitude of Low Frequency Fluctuations (fALFF), regional homogeneity (ReHo) and the eigenvector centrality. fALFF, ReHo and eigenvector centrality were extracted by using AFNI (*58*). fALFF could be interpreted as spontaneous neural activity (*59*). It was extracted by using the 3dRSFC function with a frequency bandpass to only keep frequency bands between 0.01 and 0.1 Hz. ReHo is a measure of local functional connectivity between a voxel and its nearest neighbors (*60*). It was calculated using the 3dReHo function and by considering the 27 neighborhoods of each voxel. Eigenvector centrality corresponds to a measure of general influence (*61*). It was calculated using the 3dECM function. FC was extracted using the CONN toolbox (RRID:SCR_009559) release 20.a (*62, 63*). The pre-processed images were entered to the toolbox and denoised using a standard denoising procedures (*64*) which involved removing various potential confounding effects. This included regression of the WM timeseries using 10 CompCor noise components, CSF timeseries using 5 CompCor noise components, all motion parameters along with their first derivatives (12 factors) (*65*), and scans considered outliers (below 48 factors) (*66*). Additionally, session effects and their first derivatives (8 factors) were addressed, as well as linear trends (2 factors) within each functional run (sham and ventricle stimulation). Subsequently, bandpass frequency filtering was applied to the blood-oxygen-level-dependent (BOLD) timeseries (*67*) within a range of 0.01 Hz to 0.1 Hz. CompCor (*68*) noise components for both WM and CSF were determined by calculating the mean BOLD signal and the most significant principal components orthogonal to the average BOLD signal, motion parameters, and outlier scans within each subject’s eroded segmentation masks. To examine the FC in the networks, ROI-to-ROI analysis was performed by grouping voxels into ROIs in the HPC-ICA networks (*63*). Seven functional brain networks were selected including the DMN, FPN, SN, DAN, SMN, VN, and AN (*63*). The BOLD signal time series were averaged from all voxels compromising each ROI. Fisher-transformed bivariate correlation coefficients were calculated between each pair of ROIs as reflections of connections. The intra-network FCs were calculated by averaging correlation coefficients between the ROIs within the networks.

### Statistical analysis

The statistical analyses were performed using IBM SPSS Statistics for Windows, version 28 (IBM Corporation, Armonk, NY, USA), R and SPM12. The questionnaire to evaluate the aversive effects of FUS was analysed using χ2 test.

MRI metrics analyses consisted of paired t-tests and Bayesian paired t-tests between the sham and ventricle stimulations. Whole-brain voxel-based analyses were conducted for VBM and rs-fMRI metrics since changes could rapidly spread far from the brain target. We considered as significant any clusters of size k > 25 and p _FDR-corrected_ < 0.05. All metrics were also compared between the two conditions at the region of interest (ROI) level (a cube of size 3cm^3^ around the left posterior ventricle; see Figure 3A), as well as at the functional network’s atlas level (mean connectivity within each network). Bayesian analyses results are reported as Bayes Factor (BF)10. If BF10 > 1 (H1 more likely than H0), we reported the estimated median and 95% confidence interval of the effect sizes following 10,000 iterations.

## Acknowledgments

The authors acknowledge Sarah Wilson, Louise Cowell, Andrew Cooper, Jan A Paul, and Mehri Kaviani for their MRI support in the project.

## Funding

The Academy of Medical Sciences Springboard SBF007\100077 UK (JJ)

Engineering and Physical Sciences Research Council & Medical Research Council funded NEUROMOD+ UK (JJ, MK, MALR)

Engineering and Physical Sciences Research Council EP/W004488/1 UK (MK, CA)

Engineering and Physical Sciences Research Council EP/X01925X/1 UK (MK, CA)

Engineering and Physical Sciences Research Council EP/W035057/1 UK (MK, CA)

Guangci Professorship Program of Rui Jin Hospital, Shanghai Jiao Tong University China (MK)

Medical Research Council intra-mural funding MC_UU_00030/9 UK (MALR)

## Author contributions

Conceptualization: JJ, MALR Methodology: JJ, CA, MK

Investigation: JJ, CA

Visualization: JJ, CA

Writing— JJ, CA

Writing—review & editing: JJ, CA, MK, MALR

## Data and materials availability

All data will be available prior to publication date.

