## Supplementary material for "Ventricle stimulation as a potential gold-standard control stimulation site for transcranial focused ultrasound stimulation": SI

**Supplementary information**

Supplementary Table 1

Supplementary Table 2

Supplementary Table 1

**Participant Report of Symptoms**

| Visit Date | Post time: | Investigator: |
| --- | --- | --- |
| How are you feeling overall right now? | Participant |  |
| “Right now, do you feel you have or are…? | Value  1 absent  2 mild  3 moderate  4 severe | Relation  1 unrelated  2 unlikely  3 possible  4 probable  5 definite |
| Headache |  |  |
| Unusual feeling on the skin of your head |  |  |
| Neck pain |  |  |
| Tingling |  |  |
| Itchiness |  |  |
| Difficulty paying attention |  |  |
| Unusual feelings, attitude, emotions |  |  |
| Sleepiness |  |  |
| Change in hearing |  |  |
| Nausea/stick to stomach |  |  |
| Dizziness |  |  |
| Anxious/worried/nervous |  |  |
| Forgetful |  |  |
| Difficulty with your balance |  |  |
| Other |  |  |

Supplementary Table 2

|  | Ventricle FUS | ATL FUS | χ2 | p |
| --- | --- | --- | --- | --- |
| Headache | 0 | 0 | − | − |
| Unusual feeling on the skin | 2 | 3 | 0.178 | 0.673 |
| Neck pain | 2 | 1 | 0.407 | 0.524 |
| Tingling | 3 | 3 | 0.003 | 0.953 |
| Itchiness | 0 | 0 | − | − |
| Difficulty paying attention | 3 | 2 | 0.278 | 0.598 |
| Unusual feelings, attitude, emotions | 0 | 0 | − | − |
| Sleepiness | 5 | 4 | 3.376 | 0.185 |
| Change in hearing | 1 | 0 | 1.069 | 0.301 |
| Nausea/stick to stomach | 0 | 0 | − | − |
| Dizziness | 2 | 2 | 2.188 | 0.139 |
| Anxious/worried/nervous | 0 | 0 | − | − |
| Forgetful | 0 | 0 | − | − |
| Difficulty with your balance | 1 | 1 | 0.001 | 0.974 |
| Other | 14 | 15 | 0.655 | 0.157 |
| (heard faint clicking/tapping sound during the stimulation) |  |  |  |  |

Table S2. The results of questionnaire between the ventricle and ATL sessions
